# Genipin-Crosslinked, Silane-Anchored 3D Tumor–Stroma Microtissues for High-Content On-Chip Drug Testing

**DOI:** 10.1101/2025.07.03.662913

**Authors:** Doriane Le Manach, Reza Kowsari-Esfahan, Emilia Reszczyńska, Philippe Nghe, Matthias Nees

**Affiliations:** Deptartment of Biochemistry and Molecular Biology, Medical University of Lublin, W. Chodźki 1 Street, 20-093 Lublin, Poland; Laboratoire de Biophysique et Evolution, UMR CNRS-ESPCI 8231 Chimie Biologie Innovation, PSL University, Paris, France; Department of Plant Physiology and Biophysics, Institute of Biological Sciences, Faculty of Biology and Biotechnology, Maria Curie-Skłodowska University, Akademicka 19 Street, 20-033, Lublin, Poland; FICAN West Cancer Centre Laboratory, Cancer Research Unit, Institute of Biomedicine, Turku University Hospital, University of Turku, Turku, Finland

**Keywords:** 3D tumor-stroma co-culture, silane-functionalization microfluidic devices, Genipin crosslinking, extracellular matrix stabilization, in vitro chemosensitivity assays, AI-driven high-content imaging, personalized medicine

## Abstract

Physiologically relevant 3D tumor models incorporating extracellular matrix (ECM) and cancer-associated fibroblasts (CAFs) are essential for studying tumor progression and drug resistance yet often suffer from hydrogel contraction and instability – especially in microfluidic formats, where ECM deformation hampers long-term culture and quantitative imaging. Here, we present a microfluidic tumor-stroma co-culture platform for head and neck squamous cell carcinoma (HNSCC) that overcomes these limitations through a dual-material strategy: APTES-mediated surface silanization anchors the ECM to the chip, while Genipin-based crosslinking enhances matrix stiffness without compromising cell viability. This approach stabilizes collagen-rich hydrogels for over 10 days, preserving 3D architecture, sustaining >85% viability, and supporting active proliferation. Fourier-transform infrared spectroscopy (FTIR) confirmed successful collagen crosslinking, combining covalent modification of biomaterials with improved mechanical performance. The platform further integrates AI-assisted, high-content imaging to quantify dynamic phenotypic drug responses at both single-cell and higher multicellular/tissue level resolution. Drug chemosensitivity assays, including the co-culture of tumor cells with patient-derived CAFs, demonstrated the quantitative assessment of clinically relevant chemoprotective effects. By combining biomaterial engineering with functional microfluidic design, this system enables reproducible, physiologically relevant modeling of tumor-stroma interactions, offering a scalable tool for preclinical drug screening and personalized medicine or precision oncology applications.

## 1. Introduction

Head and neck cancer (HNC) is the seventh most common cancer globally, with head and neck squamous cell carcinoma (HNSCC) accounting for the majority of cases, while other subtypes such as adenocarcinomas are much less frequent ^[1]^. Despite recent improvements in surgery, chemotherapy, and radiotherapy, the five-year survival rate for HNSCC remains close to 50%, largely due to late diagnosis, aggressive tumor biology, and limited efficacy of current treatments ^[2,3]^. Although immunotherapies have marked a significant advancement for patient outcome, their benefit is limited to only a subset of patients, posing economic and clinical challenges ^[4]^. This underscores an urgent need for more effective, accessible therapies, including pathway-specific small molecule inhibitors, personalized combination treatments, and predictive, physiologically relevant drug screening platforms for advanced and metastatic disease.

A key bottleneck in developing such therapies is the inadequacy of existing preclinical models. Traditional two-dimensional (2D) cell cultures and long-established tumor cell lines, adapted to artificial plastic substrates, lack the complex heterogeneity, spatial architecture, and microenvironmental cues that define primary tumors. Similarly, preclinical animal models, such as mouse xenografts and patient-derived xenografts (PDOs), fall short in replicating the native human tumor microenvironment (TME) ^[5,6]^. These models are often highly reductionist and fail to predict clinical drug responses accurately, resulting in costly late-stage failures in drug development.

Tumor heterogeneity adds another layer of complexity, with both genetic and non-genetic cellular diversity observed between patients and within individual tumors. This high level of tumor cell heterogeneity increases during tumor progression and contributes to therapy resistance, metastasis, and relapse ^[6]^. To better represent this biological diversity, we use early-passage tumor cell lines and patient-derived cancer-associated fibroblasts (CAFs) established directly from fresh biopsies. For example, the early-passage cell lines established from primary HNSCC biopsies preserves heterogeneity, differentiation potential, and physiological relevance of head and neck cancer cells, unlike long-term cultured lines that show massive genetic drift over time ^[7]^.

Critical components of the original TME – extracellular matrix (ECM), fibroblasts, immune cells, and endothelial cells – are typically generally absent in 2D cultures, limiting their ability to recapitulate tumor complexity and dynamic cellular crosstalk ^[8–10]^. Recent technological advances, including single-cell RNA sequencing and spatial proteomics, have revealed previously unrecognized stromal heterogeneity, identifying multiple CAF subtypes with distinct roles in tumor progression ^[11–13]^. Diverse CAFs secrete growth factors, cytokines, chemokines, and extracellular vesicles that drive immune evasion, epithelial-to-mesenchymal transition (EMT), tumor cell invasion and metastasis, therapy resistance, and ECM remodeling—contributing to tumor stiffness, fibrosis, and desmoplasia ^[14,15]^. Faithful incorporation of CAFs and tumor-specific ECM is thus critical for physiologically relevant models that more faithfully replicate the in vivo tumor architecture and potentially predict therapeutic outcomes. Collagen type I, a major ECM component in solid tumors, improves the physiological relevance of 3D culture systems due to its biocompatibility, defined chemical composition, and ability to support integrin-mediated cell adhesion, migration, and invasion of tumor cells and fibroblasts ^[16–18]^.

However, collagen-based hydrogels also introduce significant technical challenges: fibroblasts and invasive tumor cells actively remodel the collagen-rich ECM and exert traction forces that cause deformation, contraction, and possible disintegration of the hydrogel scaffold ^[19– 21]^. While this dynamic remodeling partially mimics fibrosis and desmoplasia, important features of aggressive tumors observed in vivo, it also undermines mechanical stability, reproducibility, and cell viability of the in vitro models - especially limiting their application for high-throughput drug screening and long-term assays ^[22,23]^.

To overcome these limitations, we developed a microfluidic platform (or microphysiological system) that stabilizes ex vivo tumor microtissues within a crosslinked hydrogel matrix, covalently anchored to the device surface. Our approach combines Genipin as a crosslinking agent for a composite, ECM hydrogel that is based on type I collagen (COL), hyaluronic acid (HA), and Matrigel™, with silane-based APTES surface functionalization of PDMS/glass microfluidic chips. This integrated (bio)materials strategy enhances matrix mechanical integrity and adhesion, thus effectively preventing contraction and preserving the 3D architecture of co-cultures containing patient-derived CAFs and HNSCC cell lines with varying EMT phenotypes.

Microfluidics offers precise control of microenvironmental parameters such as oxygen and nutrient gradients, mechanical forces, and chemical cues, closely mimicking in vivo tumor conditions ^[24,25]^. Additionally, microfluidic miniaturization reduces the number of primary cells needed and enables multiplexed, parallel assays for functional drug screening ^[26,27]^. Altogether, we demonstrated that our platform supports long-term, physiologically relevant 3D tumor-stroma co-cultures, enabling robust quantitative readouts for tumor invasion, matrix remodeling, and drug response via live-cell imaging, combined with automated AI-enabled analysis. Our work bridges breakthrough materials engineering with microfluidic precision, establishing a versatile tool for personalized cancer medicine, mechanistic studies of tumor-stroma interactions, and high-content functional drug screening.

## 2. Experimental Section

### 2.1 Microfluidic device fabrication

Microfluidic devices were designed in AutoCAD⍰2024 (Autodesk, San Rafael, CA, USA) and a chromium photomask prepared via Heidelberg µPG⍰101 direct writing. SU-8 features (200⍰±⍰5⍰µm height) were patterned on silicon wafers using Karl Suss MABA6 photolithography. Molds were silanized with perfluorooctyl trichlorosilane (Sigma PFOCTS) to create an anti-adhesion layer. PDMS (Sylgard⍰184) was mixed (10:1), degassed, cast (30⍰g), cured at 70⍰°C for 4⍰h, and plasma-bonded to cleaned glass slides. Inlets (⍰1mm) and outlets (3⍰mm) were punched; devices were annealed at 90⍰°C for ≥48⍰h to reestablish hydrophobicity and prevent gel leakage. The full design layout of the microfluidic chip is provided as Supplementary File S1 (Tumor_stroma_co-culture_microfluidic_chip_design - AutoCAD format) to support reproducibility and adaptation.

### 2.2 Cell culture maintenance

Human head and neck squamous cell carcinoma (HNSCC) cell lines UT-SCC-19A (laryngeal carcinoma) and UT-SCC-44 (gingival carcinoma) were provided by Turku University Central Hospital, Finland. These lines were originally derived from recurrent and metastatic HNSCC, respectively, and were selected based on their differential sensitivity to Cisplatin, as previously described in Lepikhova et al.^[7]^.

Normal human foreskin fibroblasts BJ (ATCC CRL-2522), hTERT-immortalized cancer-associated fibroblasts (PF179T), and LNCaP prostate cancer cells were obtained from the biobank of the Institute of Biomedicine, Cancer Research Unit and FICAN West Cancer Centre Laboratory, University of Turku and Turku University Hospital. Patient-derived fibroblasts were isolated from tumor biopsies according to ^[28]^ with permission and according to the guidelines of the Institutional Review and Ethical Board of the Medical University of Lublin (KB-0024/134/09/2024 and KE-0254/96/2020).

UT-SCC-19A, UT-SCC-44, BJ and patient-derived CAFs were cultured at standard growth conditions (incubator with 37 °C, 5% CO^2^) in DMEM/F-12 (Sigma, D8437), supplemented with 10% Fetal Bovine Serum (FBS, ThermoFisher, Gibco, 16140071), penicillin (100 units/ml), streptomycin (100 µg/ml), and antibiotics (Sigma, A5955). LnCAP cells were propagated in RPMI-1640 (Sigma-Aldrich, R8758).

### 2.3 3D co-culture hydrogel optimization

ECM composition was optimized in Ibidi 96-well angiogenesis plates, following the method described in Afshan et al. ^[29]^. The ECM components were diluted to final concentrations: Matrigel (Corning, 356231; 2⍰mg/mL), collagen⍰I (Corning, 354236; 0.75⍰mg/mL), and hyaluronic acid (HA, ThermoFisher Scientific, J66993-ME; 5% v/v; 10⍰mg/mL stock). HA stock was prepared by dissolving 100⍰mg HA sodium salt in 10⍰mL PBS with ions and gently stirring overnight.

### 2.4 Seeding and hydrogel polymerization in microchips

A hydrogel-cell suspension (2.5×10^6^⍰cells/mL; 1:1 cancer: fibroblast ratio) was sequentially injected into opposing gel ports to fill the central chamber. To prevent gel desiccation during polymerization, hydrogel domes were formed over the injection ports, and chips were placed in a sealed humidified chamber for 30⍰min at 37⍰°C. Following polymerization, DMEM/F-12 medium was added to the reservoirs, and the microfluidic devices were maintained at 37⍰°C with 5% CO_2_. Culture medium was refreshed every 48⍰h. Genipin (Sigma G4796) was used to crosslink and stabilize collagen-based hydrogels due to its low cytotoxicity, offering a biocompatible alternative to conventional crosslinkers such as glutaraldehyde ^[30]^.

### 2.5 Surface treatment for hydrogel adhesion

Plasma-activated PDMS and glass surfaces were silanized with 2% (v/v) (3-Aminopropyl) triethoxysilane (Sigma A3648) in MilliQ water for 2⍰h, rinsed twice in ethanol, and dried under compressed N_2_ gas to improve gel retention and to reduce fibroblast-induced gel contraction.

### 2.6 Hydrogel contraction assay

Brightfield images of gels in chips were obtained via Leica DMi1 (Leica, Microsystems) and EVOS⍰M5000 (Thermo Fisher Scientific). Gel boundaries were manually traced in Fiji (ImageJ), and contraction calculated as: *Contraction (%) = 100 × (1 - A⍰ / A_0_)*, where A_0_ and A⍰ are initial and time-point areas. Data were analyzed in GraphPad Prism⍰9.5.1.

### 2.7 Live/dead viability assay

Microchips were stained with Hoechst⍰33342 (Sigma, 382069; 5⍰µg/mL) and Propidium Iodide (PI, Sigma, 537059; ⍰1µg/mL) for 30⍰min at 37⍰°C. 3D fluorescent images were captured by Leica Thunder DMI8 (Leica Microsystems, Wetzlar, Germany) and Nikon ECLIPSE⍰Ti confocal microscopes (Nikon Instruments Inc, Melville, NY). Instant computational clearing was applied via Leica LASX. Viability was calculated using AIVIA 14.1.0 (Leica, Microsystems; Supplementary S.1.1.) as: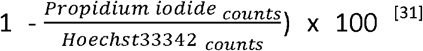. The data representation was plotted using GraphPad Prism 9.5.1 software.

### 2.8 Fourier Transform Infrared (FTIR) spectroscopy

Fourier Transform Infrared (FTIR) spectroscopy with attenuated total reflection (ATR) spectra (Nicolet⍰6700) were recorded over 4000 – 400⍰cm^−1^ at 4⍰cm^−1^ resolution (10 scans/sample), with N_2_ chamber purging. Hydrogel samples were dried on a 12-well plate then placed on an ATR diamond under N_2_ flow. The analysis was performed using OMNIC (Thermo Fisher Scientific, USA) and Grams AI software (Thermo Fisher Scientific, USA).

### 2.9 Drug Treatments

Cisplatin (Tocris) dissolved in 0.89% NaCl was administered at a range of 1 – 10⍰µM after 6⍰days of 3D culture. Treatment lasted 96⍰h, with daily drug renewal. Live/dead assay was used to determine viability; IC_50_ values were obtained via four-parameter Hill fitting in Python⍰3.11.11 (Google Colab) using log-transformed concentrations.

### 2.10 Immunofluorescence (IF) stainings

Samples were fixed in 2% PFA, permeabilized with 0.5% Triton X-100, and blocked in 3% BSA. Primary antibodies: anti-Vimentin (ab16700, Abcam/1:200), anti-pan-cytokeratin (ab7753, Abcam/1:100), EpCAM (21050-1-AP, Proteintech/1:100), N-cadherin (sc-393933 Santa Cruz/1:10), anti-TWIST1 (ab50581, Abcam/1:100), anti-ZO1 (#33-9100 Thermo Fisher Scientific/1:100) anti-αSMA (ab5694, Abcam/1:100), anti-fibronectin (ab6328 Abcam/1:100), anti-S100A4 (ab19789 Abcam/1:100), Ki67 (ab15580, Abcam/1:100) and anti-E-cadherin (#14472, Cell Signaling Technology/1:100), were incubated overnight at 4⍰°C, followed by PBS washing. Secondary antibodies (goat anti-mouse IgG-Alexa Flour Plus 555 #A32727/1:200 and anti-rabbit IgG-Alexa Flour Plus 488 #A32731/1:200; Thermo Fischer Scientific, USA) were incubated overnight at 4⍰°C. Afterwards, the samples were counterstained for actin filaments using Phalloidin Alexa Fluor 647 (#8940S, Cell Signaling Technology/1:50) and DNA/nucleus with Hoechst⍰33342(405⍰nm/1:1000, Sigma, 382069). Imaging was performed on Nikon ECLIPSE⍰Ti and Zeiss LSM980 (Airy2/Elyra7). Z-stacks captured via EC⍰Plan-Neofluar⍰10×/0.3 objective across five detection channels and processed in ZEN Blue with LSM Plus and extended-depth-of-focus methods.

### 2.11 Statistical analysis and data visualization

Graphs and statistical analyses were performed using GraphPad Prism 9.5.1 and Python 3.11.11 (Google Colaboratory). Figures were assembled and finalized using Inkscape 1.4 and Adobe Illustrator 29.0. The statistical methods used are mentioned in the captions under each figure.

## 3. Results

### 3.1 Microfluidic Device Engineering and Hydrogel Stabilization Strategy for Generating Complex Tumor–Stroma Microtissues

To recapitulate the structural and mechanical features of tumor–stroma interactions within a 3D microfluidic environment (Figures 1A), we designed a composite hydrogel system stabilized with Genipin crosslinking and covalently anchored to the chip surface by APTES. Fibroblasts, particularly CAFs, are highly contractile and remodel the surrounding ECM through force generation and biochemical signaling. Cancer cells, especially aggressive ones, can further enhance ECM degradation and remodeling, often resulting in matrix compaction and architectural collapse over time. A robust hydrogel is thus critical to maintain long-term culture stability and to preserve the tumor niche microarchitecture (Figures 1B and 1C).

**Figure 1.**
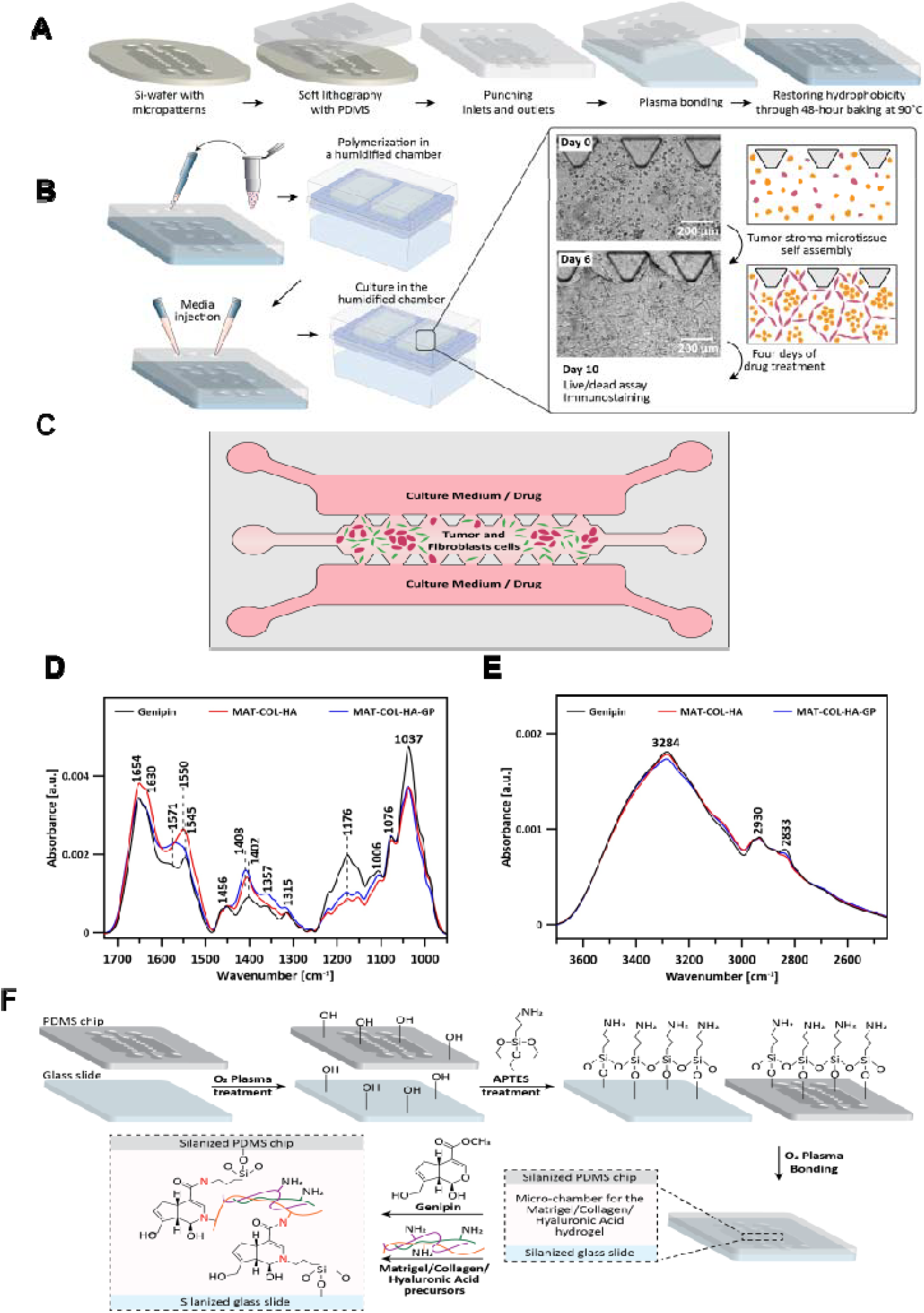
Overview of microfluidic platform development and hydrogel stabilization strategy. **(1A)** Schematic of microfluidic chip fabrication. PDMS devices were generated by soft lithography using SU-8 patterned silicon wafers, punched to create inlet/outlet ports, and irreversibly bonded to glass coverslips via oxygen plasma activation. **(1B)** Hydrogel loading and cell encapsulation workflow. Tumor cells and fibroblasts were co-seeded in a Matrigel/Collagen I/Hyaluronic Acid (Mat/COL/HA) matrix crosslinked with Genipin and injected into the central chamber. Devices were cultured for 6 days to allow microtissue formation prior to 4 days (96⍰h) of drug treatment. **(1C)** Schematic representation of the microfluidic chip layout showing the spatial distribution of tumor cells and fibroblasts within the central hydrogel chamber and adjacent side channels used for media perfusion and drug delivery. **(1D)** FTIR-ATR spectra (950–1730⍰cm^−1^) comparing Mat/COL/HA hydrogels with and without Genipin. Key shifts in amide I/II and C–O/C–N stretching regions indicate successful crosslinking and structural rearrangement. **(1E)** FTIR-ATR spectra (2450–3700⍰cm^−1^) highlighting changes in CH_2_, O–H, and N–H stretching bands. Decreases at 3284 and 2833⍰cm^−1^, along with persistent signals, suggest partial retention of functional groups relevant for subsequent surface anchoring. **(1F)** Schematic representation of the dual-surface functionalization strategy. Plasma-activated PDMS and glass were silanized using APTES to enable covalent bonding to Genipin-crosslinked hydrogels via residual hydroxyl and amine groups.

We incorporated Genipin, a naturally derived crosslinker, into a composite Matrigel–collagen type I–hyaluronic acid (Mat/COL/HA) hydrogel to reinforce its internal network via covalent interactions with primary amines, while simultaneously preserving the functional groups required for subsequent anchoring to the chip through APTES functionalization. Initial screening in 96-well plates using LNCaP/CAF co-cultures identified a concentration of 500 µM Genipin as optimal for minimizing hydrogel contraction, while being nontoxic to tumor cells or CAFs, thus fully supporting cell viability and formation of tumor-like morphologies (data not shown).

To validate matrix crosslinking and evaluate chemical functionality retention, FTIR−ATR spectroscopy was performed on Genipin alone, non-crosslinked Mat/COL/HA, and Genipin-crosslinked hydrogels after 2⍰h incubation at 37⍰°C (Figures 1D and 1E).

A distinct increase in the 1300–1450⍰cm^−1^ region (Figure 1D) corresponds to CH_2_ bending and wagging vibrations, consistent with protein backbone reorganization following crosslinking ^[32]^. The peak at 1037⍰cm^−1^, linked to –OH reorganization and potential ether or ester bond formation, also shifted slightly. Notably, a detectable signal at ⍰06⍰cm^−1^, corresponding to free Genipin, confirms that a portion remains unreacted − crucial for preserving reactive sites for chip functionalization ^[32]^.

Analysis of the Amide I (1700–1600⍰cm^−1^) and Amide II (1500–1600⍰cm^−1^) bands in FTIR-ATR revealed shifts indicative of Genipin–protein interactions (Figure 1D). The Amide II band, in particular, moved from ∼1550 to ∼1570⍰cm^−1^, possibly reflecting NH_3_^+^ symmetric bending interactions ^[33,34]^. A concurrent rise in absorbance at ∼1400⍰cm^−1^ suggests the formation of tertiary amine linkages and further crosslinking ^[32]^.

In the crosslinked formulation (Figure 1E), a reduction in the CH_2_ symmetric stretching band at 2833⍰cm^−1^ was observed, consistent with increased molecular packing and network ordering upon Genipin crosslinking^[35]^. A modest reduction in the broad band at 3284⍰cm^−1^, associated with O–H and N–H stretching, suggests partial consumption of hydroxyl and amine groups via hydrogen bonding or covalent interaction with Genipin ^[33,34]^.

Altogether, these spectral features confirm effective hydrogel crosslinking via Genipin while preserving a sufficient population of unreacted –OH and –NH_2_ groups. These groups are essential for subsequent hydrogel anchorage via oxygen plasma treatment and APTES-mediated silanization, which facilitates covalent bonding to the microfluidic chip surface (Figure 1F). This dual-functional approach ensures both mechanical reinforcement of the 3D matrix and stable spatial integration within the device over extended culture periods.

### 3.2 APTES Functionalization and Genipin Crosslinking Synergistically Enhance Hydrogel Stability and Limit Matrix Contraction in 3D Tumor–Stroma Microtissues

Following FTIR−ATR validation of Genipin-mediated crosslinking and the retention of reactive groups (Figure 1D–F), we next investigated the macroscopic impact of Genipin and APTES surface functionalization on hydrogel contraction in 3D tumor-stroma models. Preserving the hydrogel architecture over time is critical to maintain spatial, tissue-like organization and to enable long-term 3D co-culture in a mechanically stable environment.

Matrix contraction was quantified by measuring monitoring hydrogel areas over time, supplemented with two HNSCC cell lines (UT-SCC-19A and UT-SCC-44), which have been characterized as representing squamous epithelial versus partial epithelial-to-mesenchymal transition (pEMT) phenotypes, respectively (see Supplementary Figure S1). Four different experimental conditions were tested: (1) untreated control, (2) APTES surface functionalization only, (3) Genipin (500⍰µM) crosslinking only, and (4) APTES + Genipin combined (Figures 2A and 2B). On day 1, most conditions maintained the hydrogel area with minor contraction. However, UT-SCC-44 co-cultured with BJ fibroblasts showed rapid matrix collapse even after day 1, with the hydrogel area reduced to ∼13.8%, indicating pronounced contractile behavior even at early time points. This effect was markedly attenuated in Genipin-treated gels (Figure 1A).

**Figure 2.**
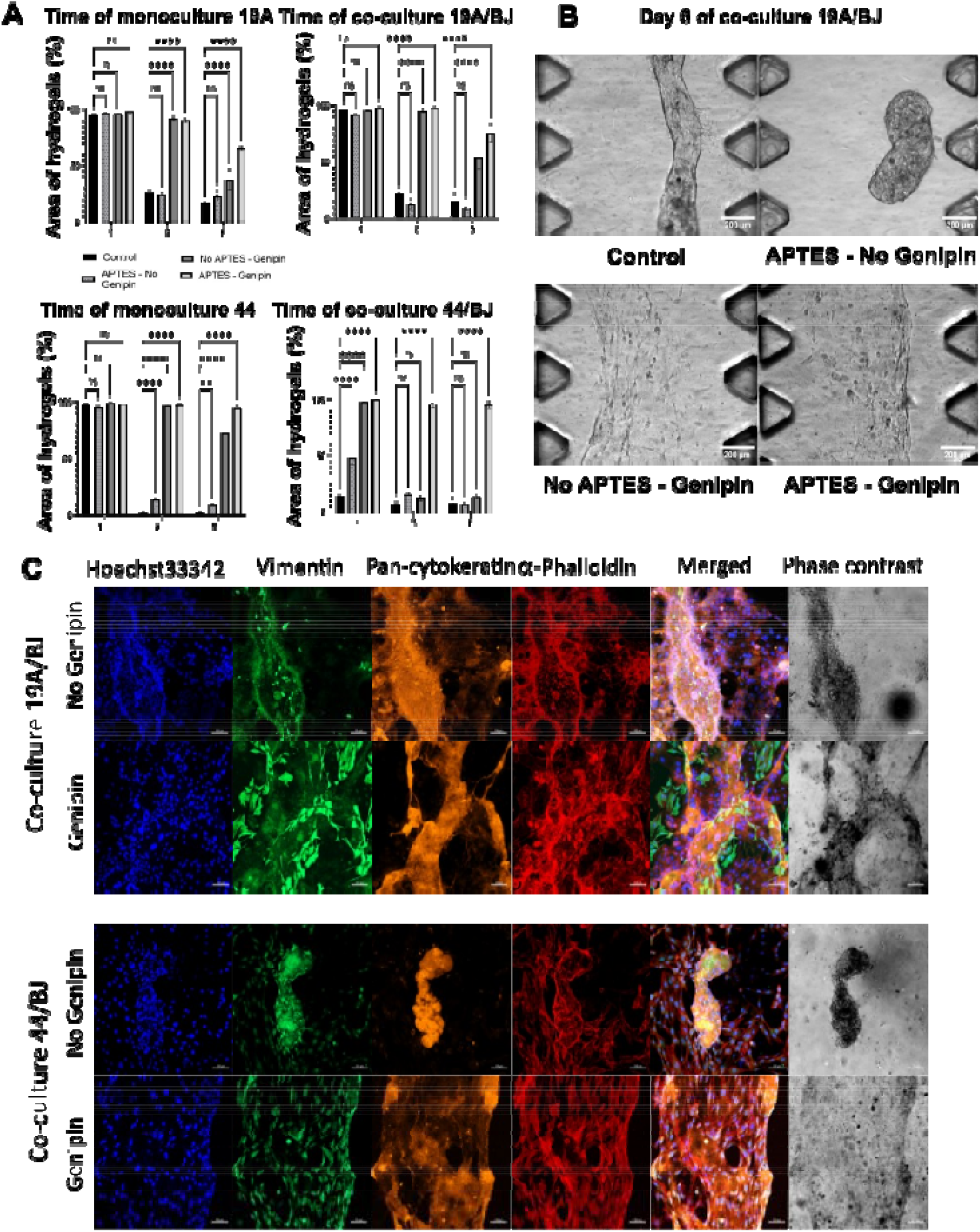
Synergistic effect of APTES functionalization and Genipin crosslinking on hydrogel stabilization in 3D tumor–stroma models. **(2A)** Quantification of hydrogel area retention over 6 days for UT-SCC-19A and UT-SCC-44 cell lines cultured in mono- or co-culture with BJ fibroblasts. Four conditions were assessed: untreated, APTES-only, Genipin-only (500rzµM), and APTES + Genipin. Data represent mean ± SD of three biological replicates. Statistical analysis was performed using two-way ANOVA with Dunnett’s post hoc test versus untreated control. (2B) Brightfield images of UT-SCC-19A/BJ co-cultures in the central chamber at day 6, comparing Genipin-crosslinked and non-crosslinked hydrogels. Scale bar: 200⍰µm. (2C) Immunofluorescence staining of UT-SCC-19A/BJ (top) and UT-SCC-44/BJ co-cultures (bottom) after 10 days, showing Vimentin (mesenchymal marker), pan-Cytokeratin (epithelial marker), F-actin (Phalloidin), and nuclei (Hoechst 33342) stainings. Z-stack projections acquired using Zeiss LSM980 with 10× objective. Scale bar: 100⍰µm.

By day 6, untreated cultures exhibited severe matrix contraction. UT-SCC-44 monocultures contracted almost completely and retained just ∼1.5% of the original hydrogel area, while co-cultures with BJ fibroblasts retained ∼7% of the original area, resulting in a dense, spherical aggregates packed with cells (Figures 2A and 2C). In contrast, Genipin-crosslinked conditions showed significant stabilization, with 72% area preserved in UT-SCC-44 monocultures and 53% in UT-SCC-19A/BJ co-cultures. Notably, the combination of Genipin crosslinking and APTES surface functionalization led to near-complete hydrogel stabilization, even observed with the highly contractile UT-SCC-44/BJ model (95% retained area). This underscores the synergistic effect of matrix reinforcement/crosslinking and anchoring to the chip (Figures 2A and 2B).

Stabilized hydrogels strongly supported the emergence of biomimetic tissue-like architecture, characteristic of squamous carcinomas (Fig 2C, top and bottom panels). Immunofluorescence (IF) staining validated the spontaneous formation of structural patterns in stabilized/cross-linked 3D co-cultures. Without crosslinking, the hydrogel rapidly collapsed into dense, disordered and feature-less clusters. In contrast, Genipin-crosslinking promoted an organized distribution of epithelial and stromal cells within the matrix, with minimal deformation of the hydrogel boundary.

In 3D cultures of the less aggressive UT-SCC-19A cells Fig 2C, top panel), Genipin cross-linking promoted the spontaneous formation of epithelial “tumor islands” within the collagen/HA matrix, surrounded by fibroblast-rich stroma. The combination of Genipin and APTES improved matrix resilience, prevented cells from escaping the matrix, and allowed localized ECM remodeling.

Remarkably, even the highly invasive UT-SCC-44/BJ co-cultures (Figure⍰2C, bottom panel) exhibited self-organized epithelial tumor islands embedded within a fibroblast-rich stroma— an architecture that this cell line typically fails to form under 2D conditions. This suggests that ECM crosslinking exerts differentiation-promoting effects and constrains tumor cell plasticity, likely through biophysical cues that recapitulate key features of the native tumor microenvironment.Together, these findings demonstrate that the combination of Genipin crosslinking, and APTES-mediated chip functionalization is critical for hydrogel stabilization, initiation and maintenance of 3D structure, promotes tissue-specific squamous differentiation, and supports spontaneous spatial organization in tumor-stroma microtissues. This integrated dual strategy enables long-term 3D co-culture and promotes the formation of complex tissue-like structures that are beyond simple organoid mono-cultures and instead represent genuine pathophysiological features observed in most solid tumors. With the help of matrix crosslinking and stabilization, we were able to establish a robust platform for mechanistic in vitro studies, such as therapeutic chemosensitivity testing in a physiologically relevant 3D environment.

### 3.3 Combined APTES Functionalization and Genipin Crosslinking Preserve Hydrogel Integrity, Enhancing Viability and Spatial Organization in 3D Tumor–Stroma Co-Cultures

Building on our demonstration that Genipin crosslinking combined with APTES surface functionalization effectively stabilizes hydrogel architecture and limits contraction (Figure 2), we aimed to show that silane functionalization of both PDMS and glass substrates enhances cell viability and in combination, more effectively prevents matrix contraction (Supplementary Figure S2). We next evaluated how this matrix stabilization impacted on long-term cell viability, proliferation, and 3D spatial organization in tumor-stroma co-cultures.

In non-crosslinked hydrogels (control and APTES-only), early matrix contraction led to scaffold collapse and dense cell clustering by Day 3, followed by increasing cell stress and death within aggregates by Day 6 and substantial escape of cells to the chip surface by Day 10 (Figure 3A). Immunofluorescence revealed that both tumor (pan-cytokeratin^+^/vimentin?) and stromal (pan-cytokeratin?/vimentin^+^) cells escaped the matrix and formed disorganized 2D layers (Figure 3B, top panel). In contrast, crosslinked hydrogels preserved matrix architecture and promoted the formation of structured tumor islands surrounded by stromal fibroblasts, mimicking tissue-like organization (Figure 3B, bottom panel).

**Figure 3:**
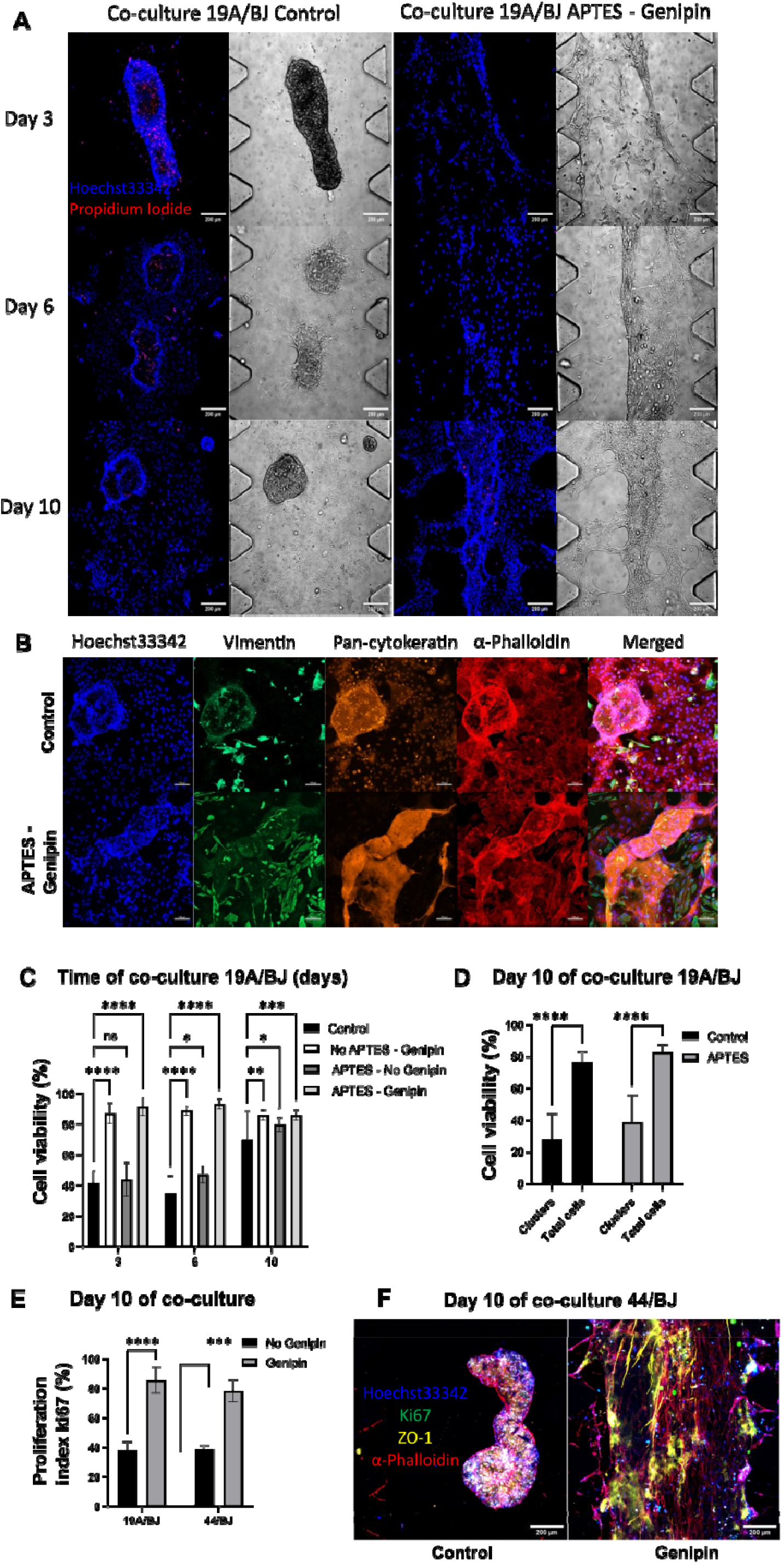
Stabilization of hydrogel architecture via dual functionalization improves spatial organization and viability of tumor–stroma co-cultures. **(3A)** Representative z-projection images of UT-SCC-19A/BJ co-cultures at Days 3, 6, and 10 cultured in either native (non-crosslinked, non-functionalized) or Genipin-crosslinked hydrogels anchored to APTES-functionalized chips. Images show phase contrast overlays with Hoechst 33342 (nuclei, blue) and Propidium Iodide (dead cells, red). Images acquired using Nikon ECLIPSE⍰Ti confocal microscope; scale bar: 200 µm. **(3B)** Immunofluorescence staining of UT-SCC-19A/BJ co-cultures at Day 10 for control and APTES/Genipin conditions, showing Vimentin (mesenchymal), Pan-cytokeratin (epithelial), Phalloidin (F-actin), and Hoechst (nuclei). Images acquired using a 10× objective (Zeiss LSM980); scale bar: 100 µm. **(3C)** Quantification of cell viability based on live/dead nuclear segmentation at Days 3, 6, and 10 across experimental conditions. **(3D)** Day 10 viability in native (non-crosslinked) hydrogels, quantifying cells retained within the matrix (“clusters”) versus the total population, including cells that migrated on the chip surface and cells that aggregated within the matrix. **(3E)** Ki67 proliferation index at Day 10 in UT-SCC-19A/BJ and UT-SCC-44/BJ co-cultures under native and crosslinked conditions. **(3F)** Z-projection IF staining of UT-SCC-44/BJ co-cultures at Day 10 in native versus crosslinked hydrogels, stained for Hoechst (nuclei, blue), Ki67 (green), ZO-1 (tight junctions, yellow), and α-phalloidin (F-actin, red). Quantification performed using automated image analysis workflows is detailed in Supplementary Methods S.1.1–S.1.2. Statistical analysis by two-way ANOVA with Tukey’s post hoc test. Data are presented as mean ± SD from two independent experiments (n = 3 biological replicates per condition). Significance: *p < 0.05, **p < 0.01, ***p < 0.001, ****p < 0.0001.

Cell viability was significantly higher in APTES–Genipin stabilized hydrogels compared to contracted conditions, where dense clusters showed increased cell death (Figure 3C). Viability analysis further distinguished matrix-resident from escaped cells, demonstrating that matrix retention strongly correlated with survival (Figure 3D). Ki67 staining confirmed active proliferation within stabilized hydrogels, while contracted cultures showed diminished proliferative capacity (Figure 3E).

High-resolution confocal imaging (Figure 3F) revealed well-organized epithelial structures in stabilized matrices, with clear epithelial polarity (ZO-1), cytoskeletal organization (α-phalloidin), and uniform nuclear distribution (Hoechst). These tissue-like features were absent in non-stabilized conditions.

Phenotypic differences between UT-SCC-19A and UT-SCC-44 were also preserved. UT-SCC-19A maintained strong epithelial traits, whereas UT-SCC-44 exhibited mesenchymal morphology and partial EMT characteristics (Supplementary Figure S1). Notably, even the aggressive UT-SCC-44 line showed spontaneous epithelial differentiation and tumor island formation within stabilized matrices, highlighting the matrix’s differentiation-promoting capacity.

Together, these results underscore the synergistic role of Genipin crosslinking and APTES anchoring in maintaining hydrogel integrity and supporting long-term viability and self-organization of 3D tumor–stroma cultures. This provides a stable and biologically relevant platform for advanced drug testing and phenotypic analysis.

### 3.4 Matrix Stabilization Modulates Single-Cell Morphometrics and Spatial Behavior in 3D Tumor–Stroma Co-Cultures

To complement the previous findings on hydrogel stability, cell viability, multicellular tissue organization and phenotype (Figure 3), we also performed a detailed quantitative morphometric analysis at the single-cell level. This approach allowed us to precisely measure how hydrogel stabilization may influence tumor and stromal cell morphology, and have an impact on spatial positioning of cells within tissue-like structures over time. Specifically, we analyzed 3D-reconstructed tumor and stromal cells cultured in native versus Genipin-crosslinked hydrogels, focusing on parameters such as cell volume, surface area-to-volume ratio (SA:V), and bounding box dimensions (see Supplementary Methods S.1.1.3 for definitions of these parameters). These quantitative metrics provide deeper insight into how matrix mechanics shape cell behavior and organization beyond qualitative observations.

### Matrix-Driven Changes in Cell Volume and SA:V Indicate Distinct Morphological States

Figures 4A and 4B show that in native, non-crosslinked hydrogels, early cell clustering and contraction leads to reduced individual cell volumes and lowered SA:V ratios by Day 3, indicative of compaction and limited nutrient diffusion. Figure 4A shows that adding Genipin (with or without APTES) stabilizes average cell sizes and volume, while these vary over a broad range in non-crosslinked gels (average during the culture of 13289 ± 6623 µm for control and 12769 ± 6137 µm for APTES only vs 19304 ± 3513 µm for APTES – Genipin). Similarly, SA:V area increases over time in native gels (from 0.26 on day 3 to 0.40 on day 10 for control and 0.29 on day 3 to 0.41 on day 10 for APTES only), while it is gradually reduced and stabilized in Genipin-crosslinked gels (from 0.46 on day 3 to 0.30 for APTES – Genipin condition; Figure 4B).

**Figure 4.**
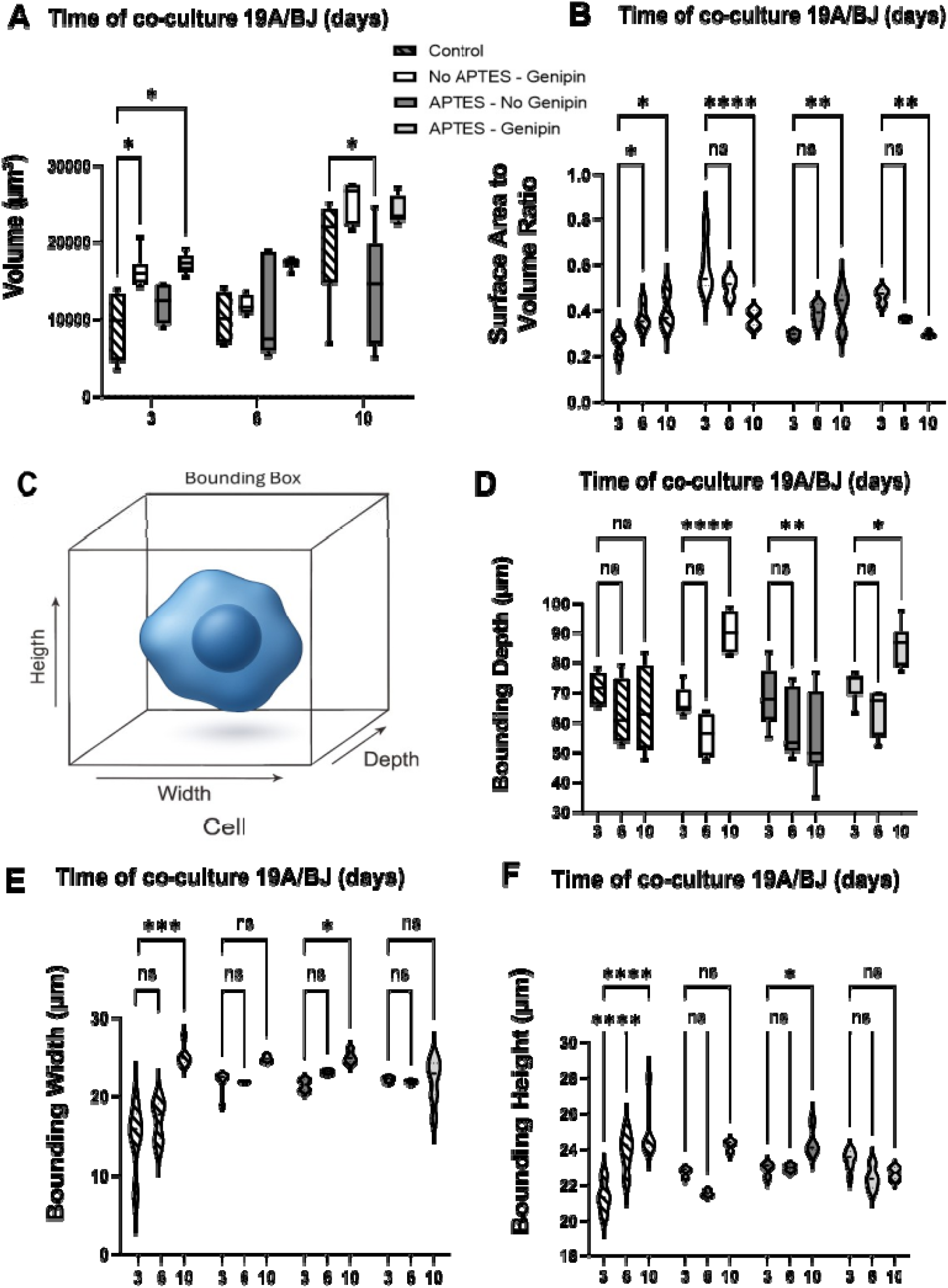
Impact of APTES functionalization and Genipin crosslinking on single-cell morphology in 3D tumor–stroma co-cultures over time. Quantitative morphometric analysis of tumor and stromal cells at Days 3, 6, and 10. (A) Cell volume (µm^3^), (B) surface area-to-volume (SA:V) ratio, (C) bounding box dimensions, and orthogonal measurements—(D) bounding depth (µm), (E) bounding width (µm), and (F) bounding height (µm)—were extracted from 3D reconstructions using Aivia software. Volume and bounding depth are shown as box plots; SA:V, bounding height, and width as violin plots. The bounding box (C) illustrates X (width), Y (height), and Z (depth) dimensions used for spatial quantification. Image analysis workflow is detailed in Supplementary Methods S.1.1. Statistical analysis was performed using two-way ANOVA with Tukey’s post hoc test. Data represents SD from three biological replicates (two chips per condition) across two independent experiments. *p < 0.05, **p < 0.01, ***p < 0.001, ****p < 0.0001.

This is partly explained by the observation that by Days 6 and 10, many cells escape the contracting matrix and instead start to adhere to the chip surface, adopting flattened but also enlarged morphologies characteristic of 2D adherent cell culture environments. In contrast, cells within Genipin-crosslinked hydrogels display a stable and consistent slow increase in volume (Figure 4A) while maintaining low SA:V ratios throughout the entire 3D culture period (Figure 4B). This suggests that matrix stabilization supports sustained volumetric growth and prevents excessive surface spreading, favoring physiological cell–cell and cell–matrix interactions within a structured 3D microenvironment.

### Bounding Box Metrics Reveal Stabilization-Dependent Constraints on Migration and Expansion

Bounding box metrics (scheme in Figure 4C) provide spatial insight into cell positioning and spreading within the hydrogel. In non-crosslinked hydrogels (control, or APTES only), the bounding depth decreases over time (from 71.1 µm on day 3 to 64.6 µm on day 10 for control; and 68.5 µm on day 3 to 60.0 µm on day 10 for APTES only), indicating vertical migration out of the matrix and onto the chip surface (Figure 4D). Concurrently, bounding width and height increase (control had a bounding width going from 20.6 µm on day 3 to 25.1 µm on day 10 and a bounding height from 21.4 µm on day 3 to 24.9 µm on day 10), reflecting lateral expansion typical of unanchored, disorganized 2D cultures (Figures 4E and 4F). Conversely, Genipin-crosslinked hydrogels exhibit increasing bounding depth (from 72.2 µm on day 3 to 85.7 µm on day 10 for APTES – Genipin condition) with stable lateral dimensions (± 23 µm of bounding height and ± 22 µm of bounding width for APTES – Genipin at different time points), consistent with deeper embedding and uniform expansion within the 3D matrix. These findings confirm that hydrogel crosslinking effectively prevents scaffold collapse and supports organized, tissue-like growth.

Together, these morphometric data extend the earlier viability and immunofluorescence results by quantitatively demonstrating how hydrogel crosslinking preserves physiological cell morphology and spatial organization. Stabilized hydrogels provide a permissive environment that supports sustained proliferation and epithelial-like organization, whereas native matrices lead to early compaction, cell stress, and disorganized migration despite rescued viability on 2D surfaces. These insights underscore the importance of matrix mechanics in shaping tumor–stroma co-culture behavior and highlight the value of crosslinked hydrogels for robust, biomimetic 3D tumor modeling and drug screening applications.

### 3.5 Genipin-Crosslinked Hydrogel Microfluidic Models Enable Drug Sensitivity Profiling in Tumor–Stroma Co-cultures

Having demonstrated that Genipin crosslinking preserves hydrogel integrity and supports long-term viability and organization of 3D microtissues, we next assessed the utility of our stabilized 3D model system for in vitro chemosensitivity testing. We focused on Cisplatin, a first-line agent in the chemotherapy of HNSCC for the past 30+ years, to examine how matrix mechanics and stromal interactions may affect therapeutic response in vitro.

### Clinical Relevance of the Cisplatin Dose Range

The range of cisplatin concentrations used (0–10⍰µM) reflected physiologically and clinically relevant exposures. Continuous infusion (40⍰mg/m^2^/day) yields patient plasma levels ∼10.7⍰µM (unbound ∼0.57⍰µM), while intraperitoneal delivery and patient variability can raise unbound levels to ∼7⍰µM. Shorter high-dose infusions (100⍰mg/m^2^ over 3⍰h) yield unbound peaks ∼6.7⍰µM ^[36–38]^. This range spans both sustained therapeutic levels and clinical peak exposures.

### Cisplatin Sensitivity Differs Between Native and Stabilized 3D Cultures

Dose–response assays with UT-SCC-19A and UT-SCC-44 in co-culture with CAFs showed reduced viability in native hydrogels starting at ≥5⍰µM Cisplatin, while Genipin crosslinking preserved cell viability across all the doses (Figure 5A). This suggests hydrogel stabilization confers some protection by maintaining structural and mechanical cues, but may also reinforce tumor–stroma interactions, which are known to support tumor cell survival and increase drug tolerance compared to monoculture conditions. In native matrices, we further distinguished responses between adherent (“bottom”) and embedded or “clustered” cells (Figure 5B): adherent cells at the bottom of our chips exhibited higher drug tolerance, which is consistent with enhanced survival signaling via integrin-mediated cell survival pathways (e.g., FAK, PI3K/AKT pathways) in 2D adherent cell cultures. In contrast, embedded cells likely faced increased mechanical stress, nutrient and oxygen limitations, and impaired polarity, reducing cell viability and increasing the overall susceptibility to cytotoxicity.

**Figure 5:**
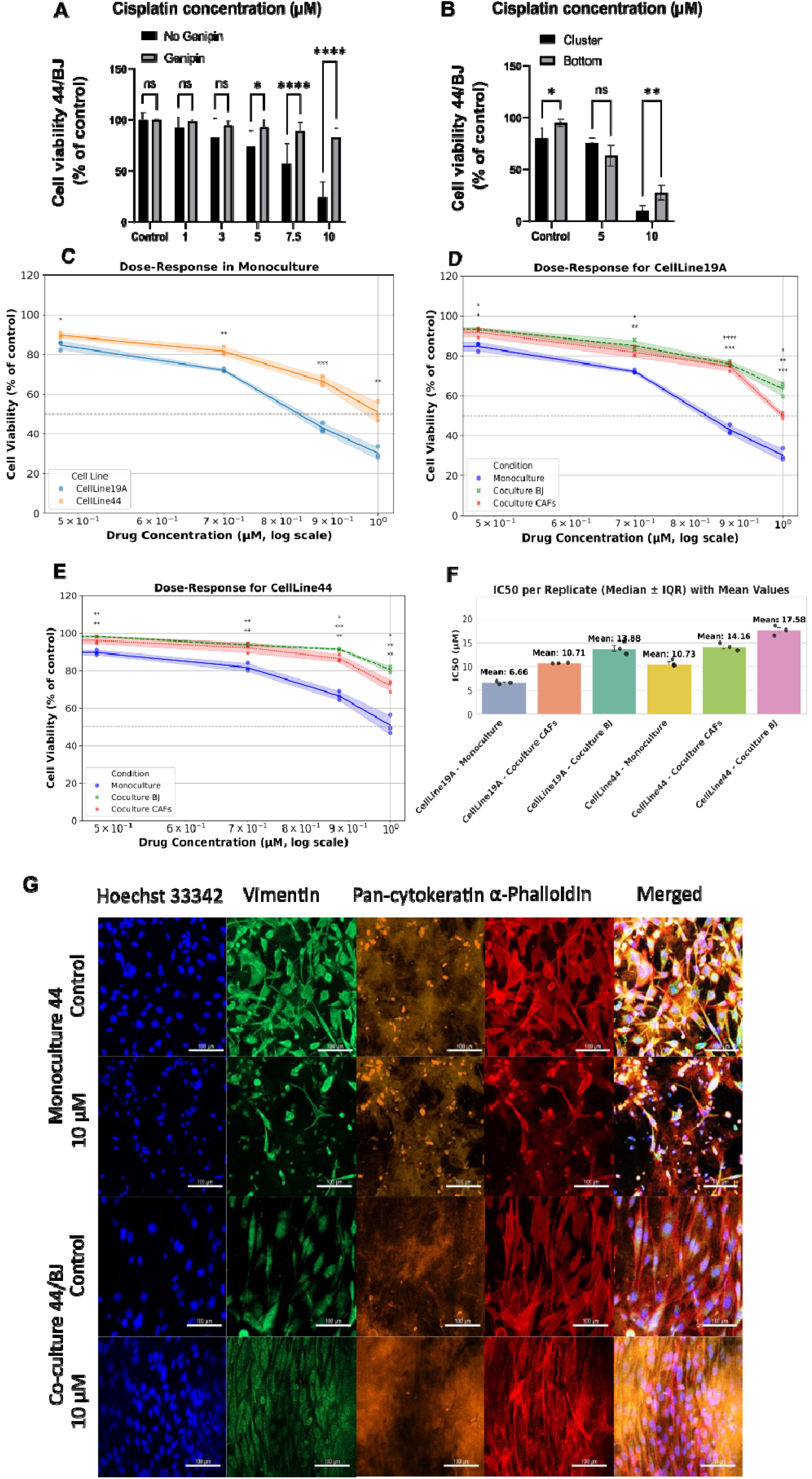
Genipin-Crosslinked Hydrogel Microfluidic Model Recapitulates Drug Sensitivity and Stromal-Mediated Chemoprotection in 3D Tumor Co-cultures. **(5A)** Cisplatin dose– response in UT-SCC-19A and UT-SCC-44 tumor–fibroblast co-cultures embedded in native (non-crosslinked) versus Genipin-crosslinked hydrogels. **(5B)** Comparison of drug sensitivity in native hydrogels between cells adhered to the chip surface and cells retained within the hydrogel matrix. **(5C-E)** Dose–response curves for UT-SCC-19A (C) and UT-SCC-44 (E) in monoculture, co-culture with BJ fibroblasts, and co-culture with patient-derived CAFs. Each dot represents a biological replicate (n = 3), with viability expressed as % of untreated control and plotted against log-transformed cisplatin concentrations. **(5F)** Summary of IC_50_ values calculated from nonlinear regression using the four-parameter Hill equation. Bars indicate the median IC_50_ (µM), with error bars representing the interquartile range (25th–75th percentile). Individual biological replicate values (n = 3) are shown as black dots; mean IC_50_ values are annotated above each bar. **(5G)** Representative immunofluorescence images of UT-SCC-44 in 3D monoculture and BJ co-culture following 4 days of Cisplatin treatment. Samples were stained for Vimentin (mesenchymal marker), Pan-cytokeratin (epithelial marker), Phalloidin (F-actin), and Hoechst 33342 (nuclei). Z-projected images acquired using a 10× objective on super-resolution microscopy. Scale bar = 100⍰µm. Statistical analysis: two-way ANOVA with Tukey’s post hoc test unless otherwise stated. *p < 0.05, **p < 0.01, ***p < 0.001, ****p < 0.0001.

### Stabilized Matrices Recapitulate EMT-Linked Drug Resistance

In crosslinked and stabilized hydrogels, UT-SCC-19A (epithelial) remained more sensitive to cisplatin than UT-SCC-44 (partial EMT), confirming that the model at least partially captures intrinsic EMT-associated chemoresistance (Figure 5C). This stratification supports its application for phenotype-specific drug screening. Notably, in previous settings, fibroblasts and CAFs have consistently shown resistance to chemotherapeutic drugs within the tested concentration range, with minimal impact on their viability at doses that effectively inhibit tumor cell growth or induce cell death ^[15,29]^. This further validates the relevance of the co-culture system for differential drug response assessment.

### Fibroblast Co-culture Confers Stromal-Mediated Chemoprotection

Fibroblast co-culture significantly increased resistance to Cisplatin in both models (Figures 5D and 5E) compared to tumor cell monoculture. In 3D monoculture (Figure 5C), the IC_50_ for cisplatin was 10.73⍰µM for UT-SCC-44 and 6.66 µM for UT-SCC-19, respectively. For UT-SCC-19A cells (Figure 5D), the IC_50_ observed for cisplatin increased from 6.66 µM observed in monoculture (blue line) to 13.88 µM in co-culture with normal fibroblasts (BJ, green line), and to 10.71 µM in co-culture with CAFs (red line). For UT-SCC-44 cells (Figure 5E), the IC_50_ rose from 10.73 µM observed in monoculture (blue line) to 17.58 µM in co-culture with normal fibroblasts (BJ, green line) and 14.16 µM with CAFs (red line). Crucially, primary patient-derived CAFs isolated from patient-derived HNSCC tissue further enhanced the clinical relevance of the model, although in these experiments, they did not enhance drug tolerance beyond the levels of BJ cells (summarized in Figure 5F). These primary CAFs maintained donor-specific heterogeneity and induced variable protective effects, underscoring the role of stromal context in modulating treatment response. Immunofluorescence confirmed preserved tumor tissue-like architecture and viability at 10⍰µM (Figure 5G), indicating significant chemoprotection mediated by the presence of stromal cells.

These findings establish hydrogel microfluidic models generated by Genipin-crosslinking of the matrix as clinically relevant platforms for drug profiling in tumor-stroma co-cultures. The system supports long-term 3D viability, preserves EMT- and stroma-linked phenotypes, and enables multiparametric readouts − including morphometric analysis (Supplementary Figure S2.3). Incorporation of patient-derived CAFs adds biological fidelity, making this a promising tool for preclinical testing and precision oncology.

## Discussion

Recreating the tumor–stroma interface in vitro remains a central challenge in cancer biology and therapeutic development. The TME is not a passive scaffold but an active driver of cancer progression, therapeutic resistance, and metastasis ^[12,13]^. In desmoplastic tumors such as HNSCC, CAFs and their mechanical remodeling of the ECM play a key role in shaping tumor behaviour ^[14,15]^. Yet most in vitro systems — including spheroids, organoids, and tumor-on-chip platforms — either completely omit the highly dynamic fibroblasts or fail to support long-term, spatially organized tumor–stroma interactions within a mechanically stable ECM ^[39–41]^.

Here, we introduce a multifunctional hydrogel-based microfluidic platform that enables sustained, spatially resolved tumor–CAF co-culture by addressing matrix contraction and hydrogel detachment—two long-standing barriers in 3D co-culture systems. Our design integrates a dual ECM stabilization strategy, combining Genipin-mediated crosslinking with APTES-based silanization, to resist CAF-induced contraction while maintaining robust hydrogel anchorage to both PDMS and glass substrates over extended durations ^[42,43]^. This allows direct tumor–CAF co-culture within a shared 3D ECM that better replicates in vivo architecture, an approach rarely utilized in vitro.

Matrix stabilization is critical in fibroblast-rich cultures, particularly under microfluidic conditions, where steep oxygen and nutrient gradients drive fibroblast activation and contractility ^[44,45]^. This commonly leads to ECM contraction, hydrogel delamination, and tissue necrosis—especially in native or non-stabilized matrices ^[46–48]^. Consequently, many co-culture systems are limited to short-term durations (<7 days), often with structural collapse and compromised viability ^[49–51]^. Our platform overcomes these limitations, supporting long-term culture (≥10 days) while preserving tissue morphology, spatial organization, and phenotype-resolved drug response.

The ECM formulation—a blend of collagen I, hyaluronic acid (HA), and Matrigel™—was optimized for bioactivity and mechanical resilience. Collagen I enables fibroblast-driven remodeling ^[52,53]^; HA contributes to increased viscoelasticity and hydration ^[54,55]^; and Matrigel™ proved essential to resist CAF-mediated contraction in collagen/HA-only matrices, likely due to its laminin-rich composition. In addition, Matrigel promoted the spontaneous self-organization of tumor and stromal cells into tissue-like structures ^[56–58]^.

To counteract matrix contraction, existing strategies typically involve trade-offs between mechanical stability, cytocompatibility, and imaging access. For example:

- Riboflavin/UVA photoinitiation offers rapid crosslinking but risks phototoxicity ^[59]^.
- Glutaraldehyde, while mechanically robust, significantly impairs fibroblast viability due to its toxicity, even at low concentrations (0.01% reduces viability by ∼70%) ^[60]^.
- PEG-based and enzymatic crosslinkers are more biocompatible but often require prolonged curing and reduce optical transparency ^[61]^.
- Mechanical constraints like microposts or confinement chambers prevent contraction but hinder perfusion and imaging ^[62]^.
- Synthetic ECMs offer tunability but often lack essential bioactivity or require functionalization to support CAF remodeling ^[63]^.

Surface coatings such as polysaccharide glue and polydopamine have been used to improve hydrogel adhesion via electrostatic or covalent interactions ^[64,65]^. However, neither method has been quantitatively validated against the strong fibroblast-mediated contraction in tumor-stroma microfluidic co-cultures. In contrast, APTES-based silanization forms covalent bonds with both the ECM scaffold and the device surface, providing durable anchorage that withstands contractile stress over extended cultures.

Our dual-stabilization strategy addresses these challenges comprehensively. Genipin, a naturally derived crosslinker, covalently bonds primary amine groups in ECM proteins while maintaining excellent cytocompatibility ^[66]^. Combined with APTES surface anchoring, this approach preserves mechanical integrity, optical clarity, and long-term viability (>85% over ≥10 days), while maintaining imaging accessibility and minimizing disruption to cell behaviour.

The stabilized hydrogel system enables non-destructive, time-resolved analysis under perfused conditions. Phenotypic metrics such as surface-area-to-volume ratios and bounding box dimensions allow real-time tracking of morphodynamic changes, cytotoxic responses, and ECM remodeling, capturing functional changes not detectable through viability assays alone.

This structural stability is essential for preserving tumor phenotype and therapeutic relevance. HNSCC microtissues maintained subtype-specific architecture, proliferation, and viability within the stabilized ECM. Native matrices, in contrast, collapsed rapidly, causing spatial disorganization. Notably, CAF co-culture significantly reduced Cisplatin sensitivity, consistent with in vivo stromal protection but rarely recapitulated in vitro ^[67]^. Subtype-specific responses were retained: the epithelial UT-SCC-19A remained more sensitive than the mesenchymal-like UT-SCC-44, reflecting its likely EMT-associated resistance ^[68]^.

Beyond pharmacologic testing, the platform has strong translational potential. Its compatibility with hydrostatic flow and low input requirements make it suitable for patient-derived samples and multiplexed assays ^[69]^. By preserving stromal dynamics and spatial organization, the system models key features of the TME increasingly recognized as modulators of drug response ^[70]^.

Looking forward, this platform offers a versatile foundation for integrating immune components and studying tumor–stroma–immune interactions. Dense or contracted ECMs are known to limit immune infiltration, particularly T cells ^[71]^. The mechanical stability and optical accessibility of our system enables quantitative studies of these transport barriers. With further development, the model could be extended to stromal-rich malignancies such as pancreatic, colorectal, or triple-negative breast cancers, and paired with high-content imaging and machine learning for morphodynamic/phenotypic analysis ^[72,73]^.

In summary, we present a materials-engineered strategy that bridges mechanical robustness and biological complexity to support long-term, spatially organized tumor–CAF co-culture in microfluidic conditions. This platform provides a structurally stable, pharmacologically informative, and translationally relevant model for cancer research, with broad applicability in precision oncology.

## Conclusion

In summary, we present a modular microfluidic platform that enables long-term 3D co-culture of tumor cells and CAFs within a mechanically stabilized, bioactive ECM. By integrating APTES-mediated surface functionalization with Genipin crosslinking, the system prevents matrix detachment and contraction − common limitations in collagen-rich microenvironments − while maintaining high viability, spatial organization, and phenotypic heterogeneity over 10 days. These advances enabled the formation of self-organized tumor microtissues with >85% viability and a 2-fold increase in Ki67 expression, supported by FTIR-ATR-confirmed crosslinking and consistent drug response profiles.

The platform supports high-content, AI-assisted morphometric analysis and multiplexed drug testing, offering a versatile toolkit for functional profiling of patient-specific tumor-stroma dynamics. From a materials perspective, this work demonstrates how rational ECM design and surface engineering can resolve key mechanical and biochemical constraints in microfluidic culture systems.

By capturing stromal-mediated drug resistance and incorporating patient-derived CAFs, this system advances a clinically relevant model for personalized oncology. Future iterations will integrate fully autologous components and on-chip analytics to support translational applications in therapy selection and disease modeling.

## Supporting information

Genipin-Crosslinked, Silane-Anchored_3D_Tumor-Stroma_Microtissues_for_High-Content_On-Chip_Drug_Testing_Supplementary_Information_Final

## Acknowledgments

We would like to thank dr Sebastian Janik from the Department of Biophysics, Instytute of Physics, Maria Curie-Skłodowska University (Lublin, Poland), led by Prof. dr hab. Wiesław I. Gruszecki, who performed super-resolution microscopy purchased with financial support from the Ministry of Science and Higher Education under grant no. 7374/IA/SP/2023, which was awarded to Prof. dr hab. Rafał Luchowski. We also thank dr hab. Justyna Widomska from the Biophysics Department of Medical University (Lublin, Poland) who has allowed us to use the Nicolet 6700 Thermo Scientific for the FTIR-ATR spectroscopic measurements. Finally, we thank Lindsey Marshall and Patrice Mascalchi from Leica Microsystems for their assistance with the use of AIVIA.

## Author contributions

Reza Kowsari-Esfahan (RKE) contributed to the microfluidic chip design and developed the dual APTES–Genipin functionalization strategy. Doriane Le Manach (DLM) led the biological implementation strategy, including tumor–stroma co-culture development, experimental validation, and data analysis. Matthias Nees (MN) contributed to experimental design refinement and translational framing. RKE developed and fabricated the microfluidic chips.

DLM performed cell culture, hydrogel experiments, immunofluorescence staining, confocal imaging, drug screening assays, viability analyses, image processing, and quantitative morphometric analysis. Emilia Reszczyńska contributed to the isolation of primary cancer-associated fibroblasts (CAFs), implementation of hyaluronic acid (HA) incorporation protocols in hydrogel preparation, and performed FTIR-ATR and materials characterization. Data interpretation was conducted by DLM. The manuscript was written by DLM with substantial support from MN and revisions from all co-authors. Project supervision was provided by DLM and MN, with co-supervision by RKE and Philippe Nghe (PN). Funding for the project was acquired by MN, PN, DLM and RKE. All authors reviewed and approved the final manuscript.

## Data Availability Statement

The data that support the findings of this study and the Python scripts are available from the corresponding authors upon reasonable request.

