## Supplementary material for "Genipin-Crosslinked, Silane-Anchored 3D Tumor–Stroma Microtissues for High-Content On-Chip Drug Testing": Genipin-Crosslinked, Silane-Anchored_3D_Tumor-Stroma_Microtissues_for_High-Content_On-Chip_Drug_Testing_Supplementary_Information_Final

**S.1. Supplementary Methods: AI-Based Image Analysis Workflow using Aivia 14.1.0 software**

**S.1.1 Cell Viability Measurements**

Cell viability was assessed using Aivia 14.1.0 software (Aivia, Leica Microsystems), a powerful AI-driven image analysis tool. The software enables pixel classification, 3D objects segmentation and quantitative analysis of volumetric image data.

**S.1.1.1 Classifier Training and Validation**

Pixel classification was trained to detect Hoechst 33342-labeled nuclei. Annotated 3D z-stacks representing varying cell densities and drug treatments were used as the training set. Pixels were categorized as either “Background” or “Want” (corresponding to the Hoechst signal). Classifier performance was evaluated by comparing detected pixels to manual annotations. Once satisfactory accuracy was achieved, the trained classifier was applied in batch mode to the entire dataset.

**S.1.1.2 3D Nuclei and Dead Cells Within Tumor Microtissues Detection**

Segmented nuclei were reconstructed using Aivia’s 3D Object Analysis (“Meshes”) module on the “Want” channel. Parameters were optimized for each experimental condition and then applied consistently. Dead cells were detected using Aivia’s Pixel Colocalization tool, using Hoechst-labelled nuclei as the mask and Propidium Iodide (PI) signal as the colocalization input. Only pixels exceeding a defined intensity threshold and showing colocalization were classified as dead cells.

**S.1.1.3 Morphological and Intensity Measurements**

Quantitative outputs (e.g., object count, size, and intensity) were extracted for each segmented 3D nucleus and dead cell object. These features were used to calculate total cell counts and viability per z-stack. Analyses were standardized across all biological replicates.

In addition, Aivia 14.1 quantified bounding depth, height, width, surface-area-to-volume ratio, and cell volume. They can be defined as following:

- Bounding Depth: The extent of the bounding box along the Z-axis, representing how far the object spans through the image stack (depth).
- Bounding Height: The extent of the bounding box along the Y-axis, indicating the object’s vertical dimension.
- Bounding Width: The extent of the bounding box along the X-axis, representing the object’s horizontal spread.
- Surface Area-to-Volume Ratio: The ratio between the object's surface area and its enclosed volume, reflecting overall shape complexity and compactness.
- Volume: The total three-dimensional space enclosed by the reconstructed cell mesh.

Graphical outputs were generated in Prism 9.5.1.

**S.1.2 IF Staining Quantifications**

Single-cell expression of epithelial, mesenchymal, and proliferative markers (e.g., Ki67) was quantified using maximum intensity projections from confocal z-stacks. Aivia 14.1.0 was used to extract fluorescence intensity values. The Corrected Total Cell Fluorescence (CTCF) was calculated by:

1. Subtracting background fluorescence (offset) from raw intensity;
2. Normalizing to ROI area (µm²);
3. Further normalizing to the number of nuclei per ROI.

This yielded CTCF per cell, accounting for both imaging conditions and cellular density.

Images were acquired on a Nikon ECLIPSE Ti confocal microscope (10× magnification, Nikon Instruments Inc., Melville, NY) with identical settings (laser power, gain, offset, HV) across all samples in each staining set.

For proliferation, the index was calculated as:

**Proliferation Index (%) = (Ki67⁺ cells / total Hoechst⁺ nuclei) × 100**

Data were visualized using Python 3.11.11 (Google Colaboratory) and GraphPad Prism 9.5.1.

**S.2 Supplementary Results**

**S.2.1 Characterization of EMT Marker Expression in HNSCC Cell Lines**

To establish a robust and physiologically relevant platform for long-term drug screening, we selected two HNSCC cell lines exhibiting distinct epithelial and mesenchymal characteristics. This enabled us to capture representative tumor phenotypic diversity within the functionalized tumor-stroma microtissues on chip and evaluate the system’s ability to sustain viability and stability of highly invasive, dynamic cells with mesenchymal traits over extended culture.

UT-SCC-44 displayed an elongated morphology and gene expression profile consistent with epithelial-to-mesenchymal transition (EMT) ^[1]^, including the partial loss of epithelial features such as reduced cell–cell adhesion and polarity, alongside increased motility and invasiveness ^[2]^. We quantified this EMT status via immunofluorescence staining of key markers and Corrected Total Cell Fluorescence (CTCF) analysis (Figure S1B) ^[3]^. Compared to UT-SCC-19A, UT-SCC-44 showed significantly higher expression of mesenchymal markers (Vimentin, TWIST1, α-SMA, S100A4, Fibronectin) and retained expression of classical epithelial markers (E-cadherin, EpCAM, pan-cytokeratin, ZO-1), indicating a partial EMT (pEMT) state (Table S1) ^[4,5]^. This hybrid phenotype is linked to increased tumor cell plasticity and metastatic potential, features that our 3D microfluidic model successfully supports.

By sustaining such heterogeneous cellular phenotypes, the platform faithfully mimics in vivo tumor complexity, providing a relevant environment for studying tumor progression and drug responses.

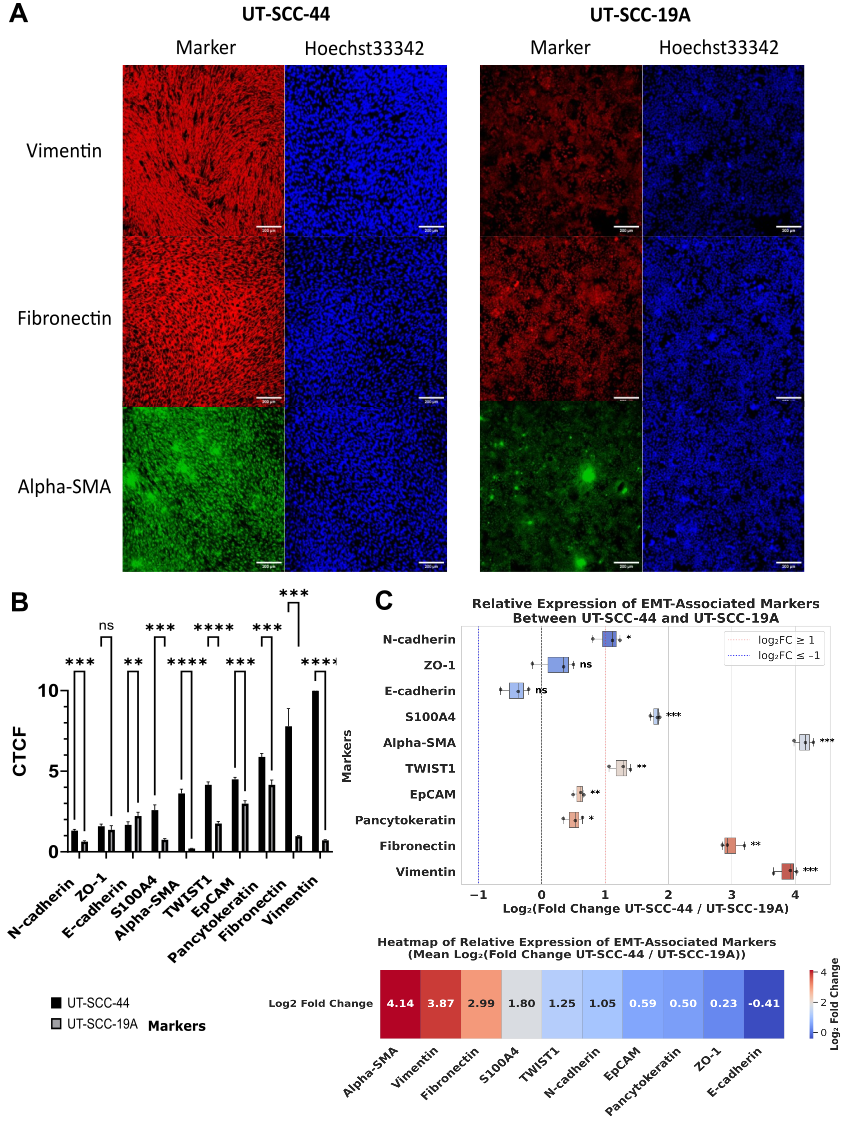
***Figure S1:*** ***Differential Expression of EMT Markers in UT-SCC-44 vs. UT-SCC-19A Assessed by Immunofluorescence and Quantitative Image Analysis. (S1A)*** *Representative immunofluorescence (IF) images of the three most upregulated epithelial–mesenchymal transition (EMT) markers in UT-SCC-44 versus UT-SCC-19A cells. Stainings were acquired at 10× magnification; scale bar: 200 μm.* ***(S1B)*** *Quantification of corrected total cell fluorescence (CTCF) for each marker based on three biological replicates.* ***(S1C)*** *Log₂ fold change in EMT marker expression between UT-SCC-44 and UT-SCC-19A. Data are presented as mean ± SD. Statistical significance was determined using one-way ANOVA with Tukey’s post hoc test: **p < 0.01, ***p < 0.001, ****p < 0.0001.*

**Table S1.** Functions and their associated expressions in EMT process of the markers tested by IF stainings

| **Markers** | **Functions** | **Expressions in EMT** |
| --- | --- | --- |
| E-cadherin | Epithelial marker, essential for epithelial cell-cell adhesion | Low |
| Vimentin | Type III intermediate filament protein, mesenchymal marker | High |
| EpCAM | Epithelial marker, involved in cell adhesion | Low |
| Pan-cytokeratin | Epithelial marker, detects a large panel of epthelial intermediate filament proteins | Low |
| TWIST1 | Transcription factor, promoting EMT | High |
| N-cadherin | Marker for loss of epithelial integrity, increasing cell motility, and facilitating invasive and metastatic behavior | High |
| α-SMA | Marker for mesenchymal cells and myofibroblasts, promotes cancer progression and metastasis | High |
| ZO-1 | Marker for epithelial cell integrity and cell-cell adhesion | Low |
| S100A4 | Fibroblast-specific protein 1 (FSP1), mesenchymal, associated with cell motility, invasiveness and metastasis | High |
| Fibronectin | Mesenchymal ECM protein, promotes cell adhesion, migration and tissue repair | High |

**S.2.2 APTES Surface Functionalization of PDMS–Glass Enhances Hydrogel Anchoring in Microfluidic 3D Tumor–Stroma Models**

Building on previous results demonstrating that Genipin crosslinking and APTES-treated glass surfaces enhanced hydrogel stabilization (Figure 2), we next assessed whether extending APTES silanization to both the glass and PDMS surfaces of the microfluidic chip would further reinforce ECM anchorage. This “full anchorage” approach was hypothesized to mechanically tether the hydrogel to all four channel walls, thereby minimizing contraction and improving long-term culture stability.

Two configurations were compared: (1) standard anchorage using APTES-functionalized glass only and (2) full anchorage involving APTES treatment of both the glass bottom and PDMS sidewalls. Quantitative image analysis showed that full anchorage significantly limited hydrogel contraction at both Day 7 and Day 10 (Figure S2A), as well as improved structural integrity in Z-projected images (Figure S2C). These findings suggest that lateral anchorage via PDMS functionalization reduces matrix detachment and compaction under cellular tension, preserving microtissue architecture.

In parallel, full anchorage was associated with higher cell viability (Figure S2B), further indicating that mechanical stabilization of the matrix benefits cell survival. This is likely due to reduced mechanical strain and more consistent maintenance of the 3D microenvironment, which supports epithelial–stromal interactions and tissue homeostasis. Representative images confirmed enhanced retention of hydrogel structure and reduced cell death in the dual-treated condition (Figure S2C).

Together, these results establish dual-surface APTES functionalization as a practical and effective strategy to improve hydrogel anchorage in microfluidic systems. By mitigating matrix deformation and enhancing viability, this method strengthens the reliability and physiological relevance of the 3D tumor–stroma co-culture platform for extended culture periods and high-content functional assays.

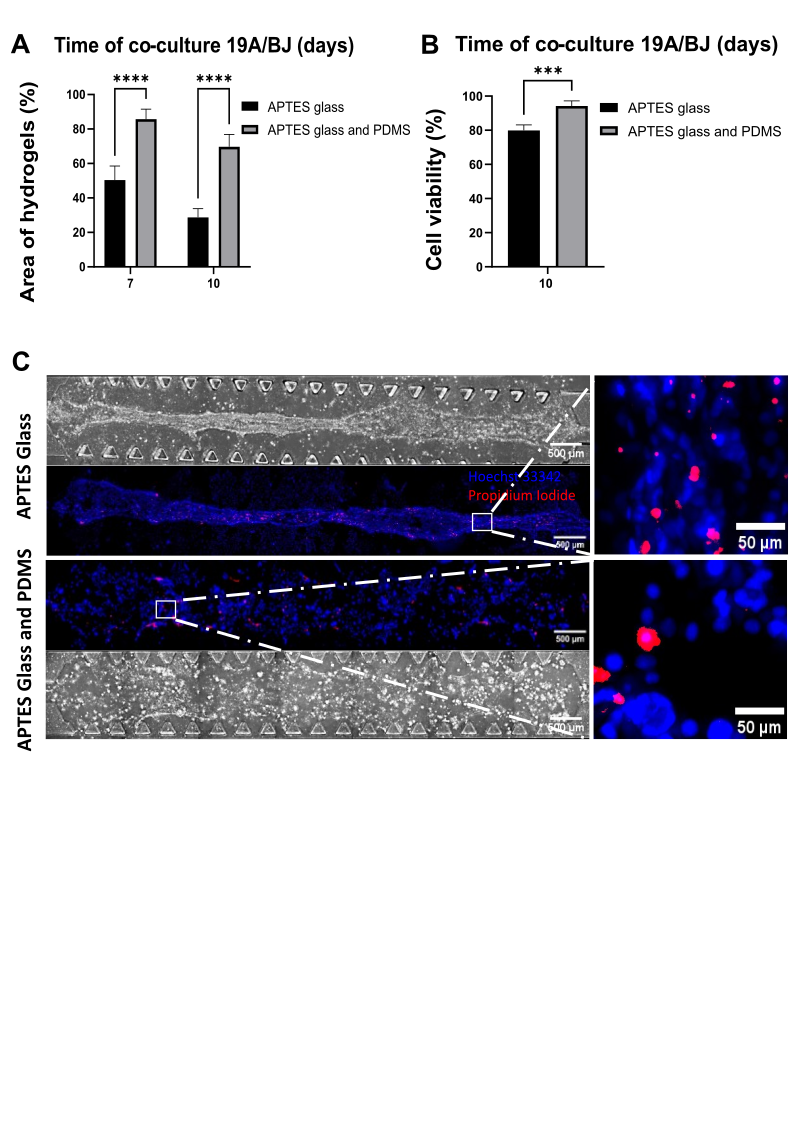
***Figure S2. Impact of Glass and/or PDMS Functionalization by APTES on hydrogel area, viability, and morphology in 3D co-cultures of UT-SCC-19A cells with BJ fibroblasts.*** ***(S2A)*** *Quantification of hydrogel area at Days 7 and 10 in UT-SCC-19A/BJ co-cultures, comparing APTES functionalization of the glass bottom versus both PDMS and glass surfaces.* ***(S2B)*** *Corresponding cell viability measurements under the same conditions at Days 7 and 10.* ***(S2C)*** *Representative confocal Z-projections at Day 10, showing brightfield images alongside Hoechst 33342 (live nuclei) and propidium iodide (dead cells) staining. Scale bars: 500 µm (overview), 50 µm (magnified region). Statistical analyses: t* *wo-way ANOVA with Sidak’s multiple comparisons test (A); unpaired t-tests (B, C); significance indicated as ***p < 0.001, ****p < 0.0001. Data represent two independent experiments with three biological replicates per condition (n = 3), analysed by full-chip imaging to ensure robustness.*

**S.2.3 Morphological Parameters Reflect Drug-Induced Stress and Resistance**

To complement the drug response findings in Figure 5, we analysed tumor cell morphology to determine whether structural changes could serve as early indicators of cytotoxic stress. Using AIVIA-based 3D segmentation, we quantified the surface area-to-volume (SA:V) ratio of individual tumor cells from UT-SCC-44 monocultures and co-cultures across increasing Cisplatin concentrations (Figure S3). In monocultures, SA:V significantly progressively decreased from 3 μM onwards (0.4257 for SA:V control vs 0.3945 for SA:V at 3 μM), consistent with morphological hallmarks of cytotoxicity or early apoptosis. By contrast, in co-cultures, SA:V values remained stable, with only a modest significant reduction at 10 μM, suggesting stromal protection and preserved cellular architecture.

These results underscore the added value of morphometric profiling as a quantitative, structure-based readout to complement viability metrics in functional drug testing. Together with the viability and proliferation data, this analysis highlights the capacity of the platform to support multi-parametric drug response profiling in a physiologically relevant 3D context.

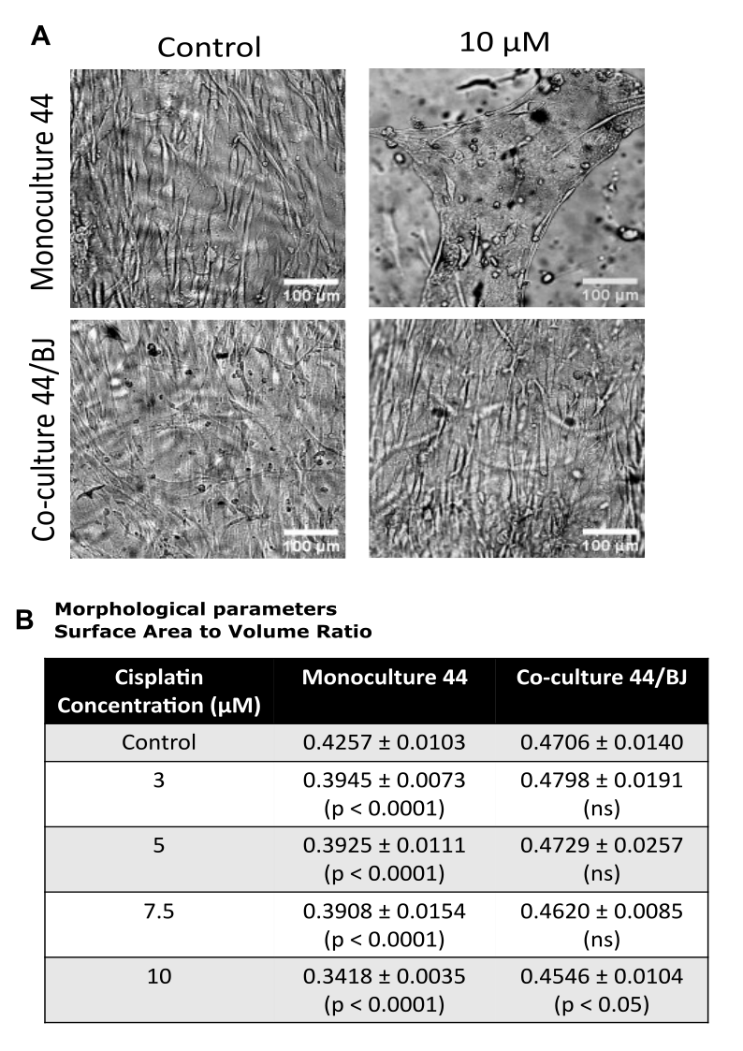

***Figure S3. Stromal co-culture modulates tumor cell morphology and drug response under Cisplatin treatment. (S3A)*** *Brightfield images of UT-SCC-44 monocultures and co-cultures ±10 μM Cisplatin, illustrating differential survival. Scale bar: 100 μm.* ***(S3B)*** *Quantification of surface area-to-volume ratio (SA:V) of individual tumor cells across Cisplatin concentrations in mono- and co-culture conditions. Approximately 700–800 cells per condition were analysed. Data represent mean ± SEM from a representative experiment. Statistical analysis: Two-way ANOVA with Sidak’s multiple comparisons test for (A–B); unpaired t-tests for comparisons at each concentration; Kruskal–Wallis test with Dunn’s post hoc correction for SA:V distributions.*

**Supplementary Files**

**File S1.** AutoCAD (.dxf) file of the microfluidic chip layout used for device fabrication. Includes dimensions, channel geometry, and inlet/outlet positioning for PDMS molding.
